# Nutrients: trace metals, micronutrients, oestrogen and B-vitamin content of Osun River: A River that runs southwestern Nigeria into the Atlantic Gulf of Guinea

**DOI:** 10.1101/2020.08.20.259010

**Authors:** N.B. Afolabi-Balogun, O.A. Oni-Babalola, I.I. Adeleke, F.A. Oseni, R.H. Bello, M. Bashir, B.A Raji

## Abstract

Osun-Osogbo Grove has a long history of healing and therapeutic claims by adherent believers, in spite of advancement in medicine. Scientists made attempts at investigating the biodiversity of the Grove, till date, there has not been convergence point between science and these indigenous beliefs. This study identified the presence of therapeutic agents in the water of Osun-Osogbo River, paying attention to at least six parameters; vitamin, phosphate, nitrate, amino acid, hormone and trace metal. Water samples were taken from two different sites during pre, during and post raining sessions (April 2017 - September 2019) were analysed using High Performance Liquid Chromatography (HPLC), Gas Chromatography Mass Spectroscopy (GC-MS) and Atomic Absorption Spectrometer (AAS). Trace metal analysis revealed an average of 0.009-0.079 mg/Kg Zinc from site one and lower in site two. The mean value of manganese at both sites was virtually the same at 0.018-0.313 mg/kg, aluminum content was 0.045-0.179mg/Kg at site one, 0.050-0.192mg/kg at site two, cobalt was 0.024 mg/kg at site one, 0.026 mg/kg at site two while nickel was 0.006 mg/kg and 0.004 mg/kg for site one and two respectively. HPLC analysis shows mean Methionine content at both sites is higher than the FDA (56.6 ug/mL); site one had 74.41 ug/mL while site two had 57.11 ug/mL The mean values of two water-soluble vitamins; Thiamine (B1) was 3.758 mg/Kg and 2.355 mg/Kg while Pyridoxine (B6) was 0.108 mg/Kg and 0.072 mg/Kg at site one and two. GCMS analysis of steroidal content revealed values below lowest observed effect level (LOEL), testosterone (4.8 ng/L) and estrogen (2.4 ng/L) were still elevated while ethinylestradiol and estriol were ≥1.5 ng/L. Summarily, site one the major part for spiritual activities showed higher essential nutrient contents than site two which support the enrichment and potential therapeutic properties of the Osun river water. However, further scientific research is required to ensure that these therapeutic potentials supersede the toxicological effect.

## 1. Introduction

Water is an essential nutrient for all known forms of life, it is also the medium by which fluid and electrolyte homeostasis is maintained in humans. The presence of liquid water on the surface is one of the key characteristics that make the Earth a unique planet, and it is usually considered to be critical for a planet to be habitable (Kasting, 2003). The composition of water varies widely with local geological conditions. Generally, all water bodies be it groundwater, surface water or any other forms have other chemical components dissolved in it. Water contains small amounts of gases, minerals and organic matter of natural origin (Sadgir and Vamanrao, 2003). Since water acquires its constituents from contact with rocks, soil and the environment, it is natural therefore to detect inorganic chemicals in drinking water that are occurring naturally. Drinking water supplies may contain some of these essential minerals naturally or through deliberate or incidental addition. Water supplies are highly variable in their mineral contents, while some contribute appreciable amounts of certain minerals either due to natural conditions (E.g., Ca, Mg, Se, F, Zn), intentional additions Fluorine (F), or leaching from piping copper (Cu), most provide lesser amounts of nutritionally/essential minerals. (Olivares and UauyZ, 2000). Prominent amidst these constituents are micronutrients, which are nutrients, required by organisms throughout life in small quantities to orchestrate a range of physiological functions. These may include; vitamins, amino acids, minerals as well as metals of enzymatic importance. Micronutrients are vital for the proper functioning of all the body systems, enabling the body to produce enzymes, hormones, and other substances essential for proper growth and development. Although require in minute quantities, absence or decrease in quantities below body requirements may have consequences ranging from mild to severe (Hannah Ritchie 2017).

Nutrients and other life-sustaining elements delivered to water body systems are a major determinant of the functioning of such water body. Thus, the Oṣun River that flows southwards through central Yoruba land in southwestern Nigeria into the Lagos Lagoon and the Atlantic Gulf of Guinea is expected to be rich in terms of micronutrients considering the climatic, and vegetation of its location. It is one of the several rivers ascribed in local mythology to have been, women who turned into flowing waters after some traumatic event frightened or angered them. Its main source is from Ilase, Osun State 7°44’34.5”N 4°39’17.6”E. The river is named after the Oṣun or Oshun, one of the most popular and venerated Orishas (Murrell, 2009). Oṣun is one of the river goddesses in Yoruba land, she is believed by devotees to provide their needs, which ranges but not limited to ability to give barren women babies and change the lives of many other people. Hence, the river water is known for its ability to cure ailments/illnesses and solve fertility problems.

There is growing concern on environmental issues and the supernatural power associated with Osun river water, mainly due to the fact that most report are unpublished and there are limited scientific information on both river constituent and its effects on believer who comes from far and near during the festival to collect water for various reasons. Olajiire and Imaokparia (2000 and 2001) reported the metals concentrations as well as inorganic nutrients of the Osun River, however, the concentrations of micronutrient in the river are uncertain. In addition to inorganic nutrients and trace metals, evaluation of the soluble B-vitamins (thiamin B1, riboflavin B2, pyridoxine B6, biotin B7 and cobalamin B12), hormone and the amino acid methionine is essential to give idea of the metabolic potential of the river water. Thus, this study is set out to identify, if any the presence of therapeutic agents/metabolic cofactors in the Osun River water.

## 2. Method

### Sampling Area

This was conducted within the Osun - Osogbo Sacred grove which is located along the bank of Osun River in Osogbo capital city of Osun State, South Western Nigeria. It is located on a latitude of 7°45’05.9”N and longitude of 4°33’03.9”E as indicated (figure1), 250km north of Lagos, land size of 75 hectares and about 350 metres above sea level. The Groves houses hundred shrines, sculptures and world heritage site (National Commission for Museums and Monuments (NCMM), 2005 and Osegale, *et al*. 2014).

**Figure 1:**
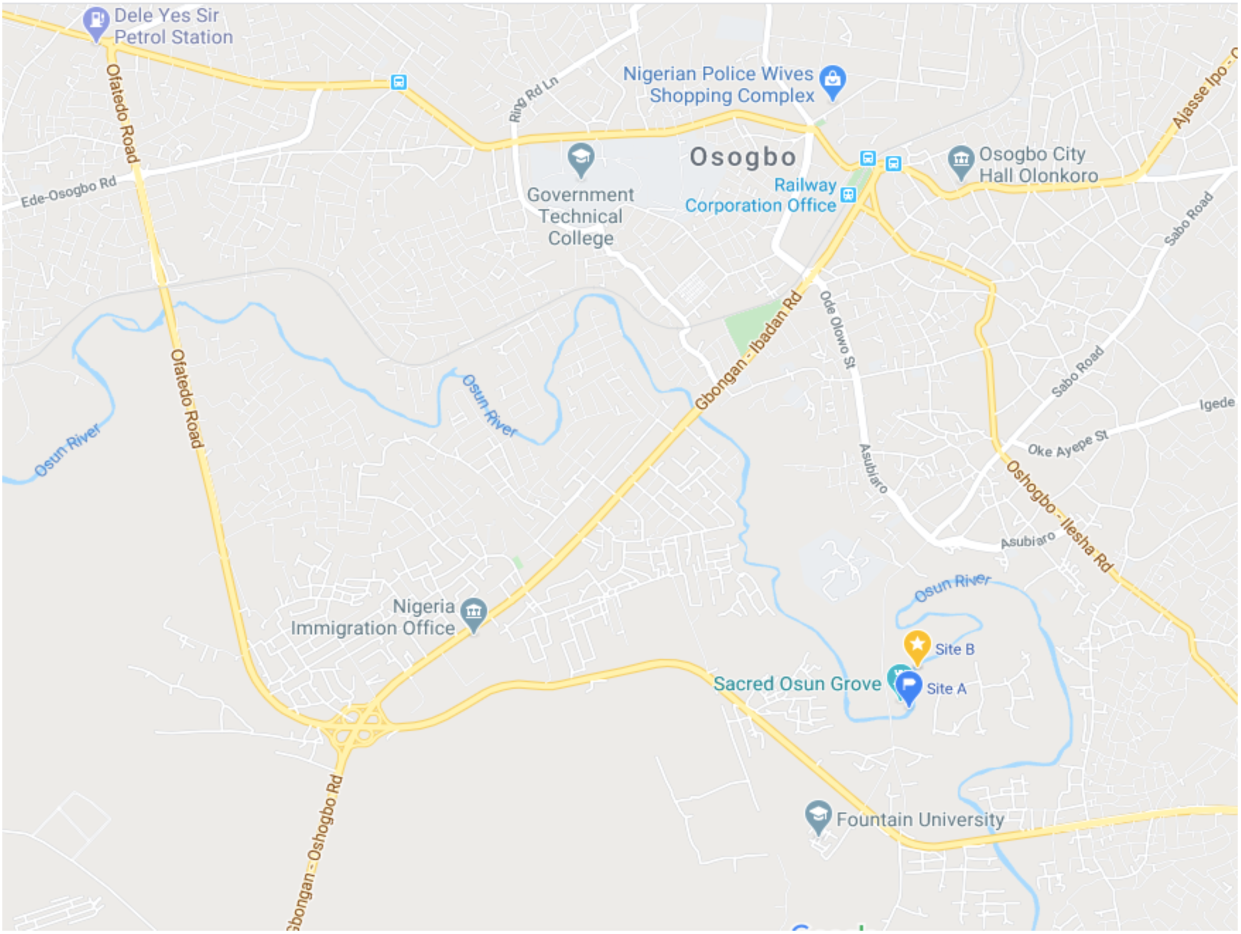
Graphical location of sampling location along the Osun River Path. Source: Map Data@2020 (maps.google.com)

**Plate 1:** A screenshot of trace metal analysis for both sites.

| Date Time | Label | Element Label (nm) | Conc | %RSD | Unadjusted Conc | Intensity | %RSD |
| --- | --- | --- | --- | --- | --- | --- | --- |
| 6/6/2017 13:21:19 | Site 2 | Al (396.152 nm) | 0.192 (ppm) | 1.96 | 0.192 (ppm) | 4834.446 | 2.04 |
| 6/6/2017 13:07:31 | Site 1 | Zn (213.857 nm) | 0.079 (ppm) | 3.84 | 0.079 (ppm) | 1235.343 | 3.49 |
| 6/6/2017 13:07:31 | Site 1 | Cd (228.802 nm) | 0.000 u (ppm) | > 100.00 | 0.000 (ppm) | -2.164 | > 100.00 |
| 6/6/2017 13:07:31 | Site 1 | Cu (324.754 nm) | -0.261 u (ppm) | 1.34 | -0.261 (ppm) | -13801.281 | 1.34 |
| 6/6/2017 13:07:31 | Site 1 | Ni (352.454 nm) | 0.006 (ppm) | 3.98 | 0.006 (ppm) | 118.935 | 3.95 |
| 6/6/2017 13:07:31 | Site 1 | Co (340.512 nm) | 0.024 (ppm) | 2.02 | 0.024 (ppm) | 291.971 | 1.99 |
| 6/6/2017 13:07:31 | Site 1 | Pb (405.781 nm) | -0.001 u (ppm) | > 100.00 | -0.001 (ppm) | -2.796 | > 100.00 |
| 6/6/2017 13:07:31 | Site 1 | Mn (403.076 nm) | 0.313 (ppm) | 0.56 | 0.313 (ppm) | 11614.817 | 0.51 |
| 6/6/2017 13:07:31 | Site 1 | Cr (425.433 nm) | 0.000 (ppm) | > 100.00 | 0.000 (ppm) | 2.089 | > 100.00 |
| 6/6/2017 13:07:31 | Site 1 | Al (396.152 nm) | 0.179 (ppm) | 2.32 | 0.179 (ppm) | 4508.113 | 2.41 |
| 6/6/2017 13:21:19 | Site 2 | Zn (213.857 nm) | -0.015 u (ppm) | 2.28 | -0.015 (ppm) | -254.419 | 2.28 |
| 6/6/2017 13:21:19 | Site 2 | Cd (228.802 nm) | -0.003 u (ppm) | 60.49 | -0.003 (ppm) | -35.049 | 60.49 |
| 6/6/2017 13:21:19 | Site 2 | Cu (324.754 nm) | -0.222 u (ppm) | 0.35 | -0.222 (ppm) | -11742.247 | 0.35 |
| 6/6/2017 13:21:19 | Site 2 | Ni (352.454 nm) | 0.004 (ppm) | 2.19 | 0.004 (ppm) | 79.570 | 2.18 |
| 6/6/2017 13:21:19 | Site 2 | Co (340.512 nm) | 0.026 (ppm) | 1.55 | 0.026 (ppm) | 317.639 | 1.52 |
| 6/6/2017 13:21:19 | Site 2 | Pb (405.781 nm) | -0.006 u (ppm) | 10.44 | -0.006 (ppm) | -23.467 | 10.44 |
| 6/6/2017 13:21:19 | Site 2 | Mn (403.076 nm) | 0.313 (ppm) | 0.14 | 0.313 (ppm) | 11616.365 | 0.12 |
| 6/6/2017 13:21:19 | Site 2 | Cr (425.433 nm) | -0.002 u (ppm) | 9.59 | -0.002 (ppm) | -76.362 | 9.59 |

**Figure 1:**
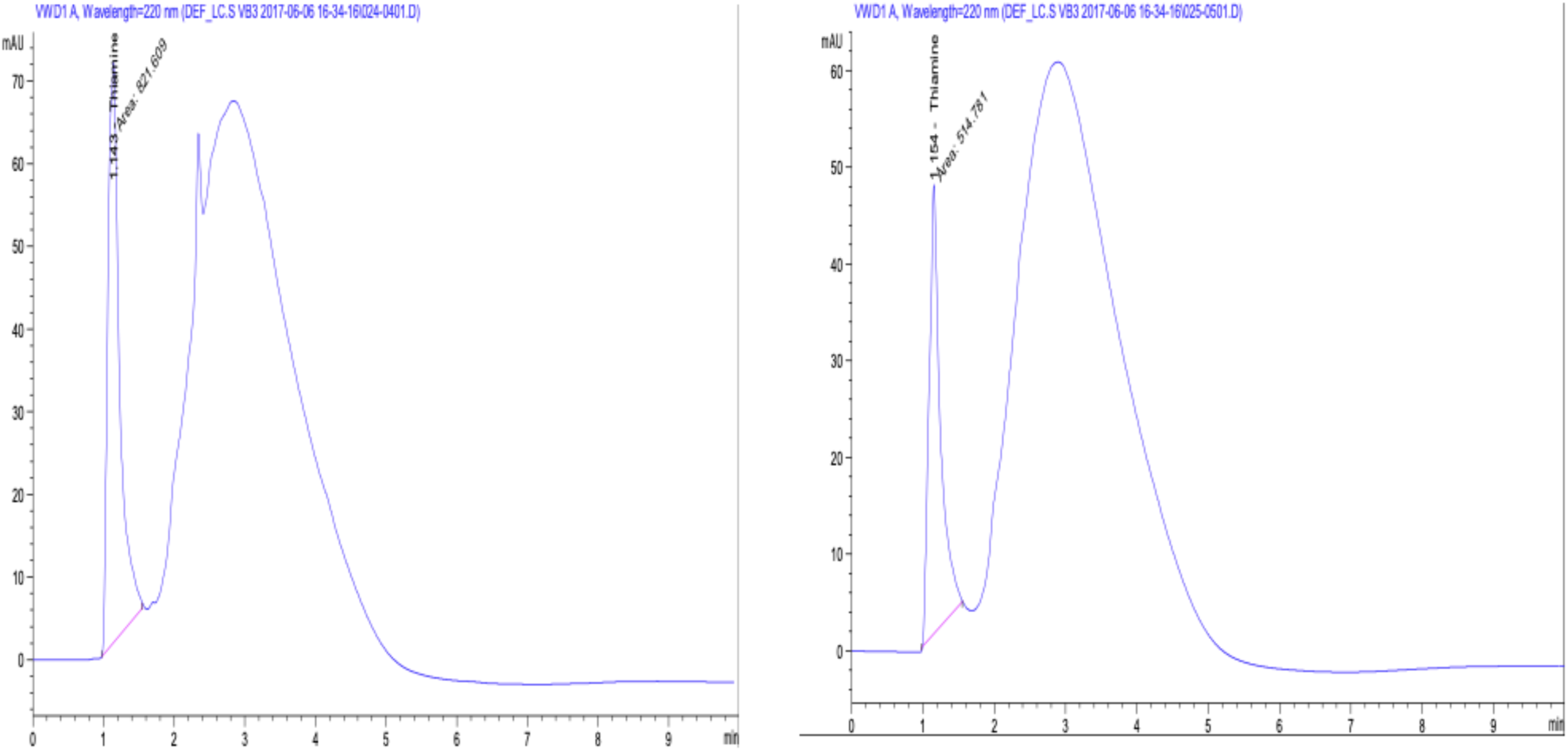
HPLC Spectra of Thiamine Content of Site 1(a) and 2 (b)

**Figure 2:**
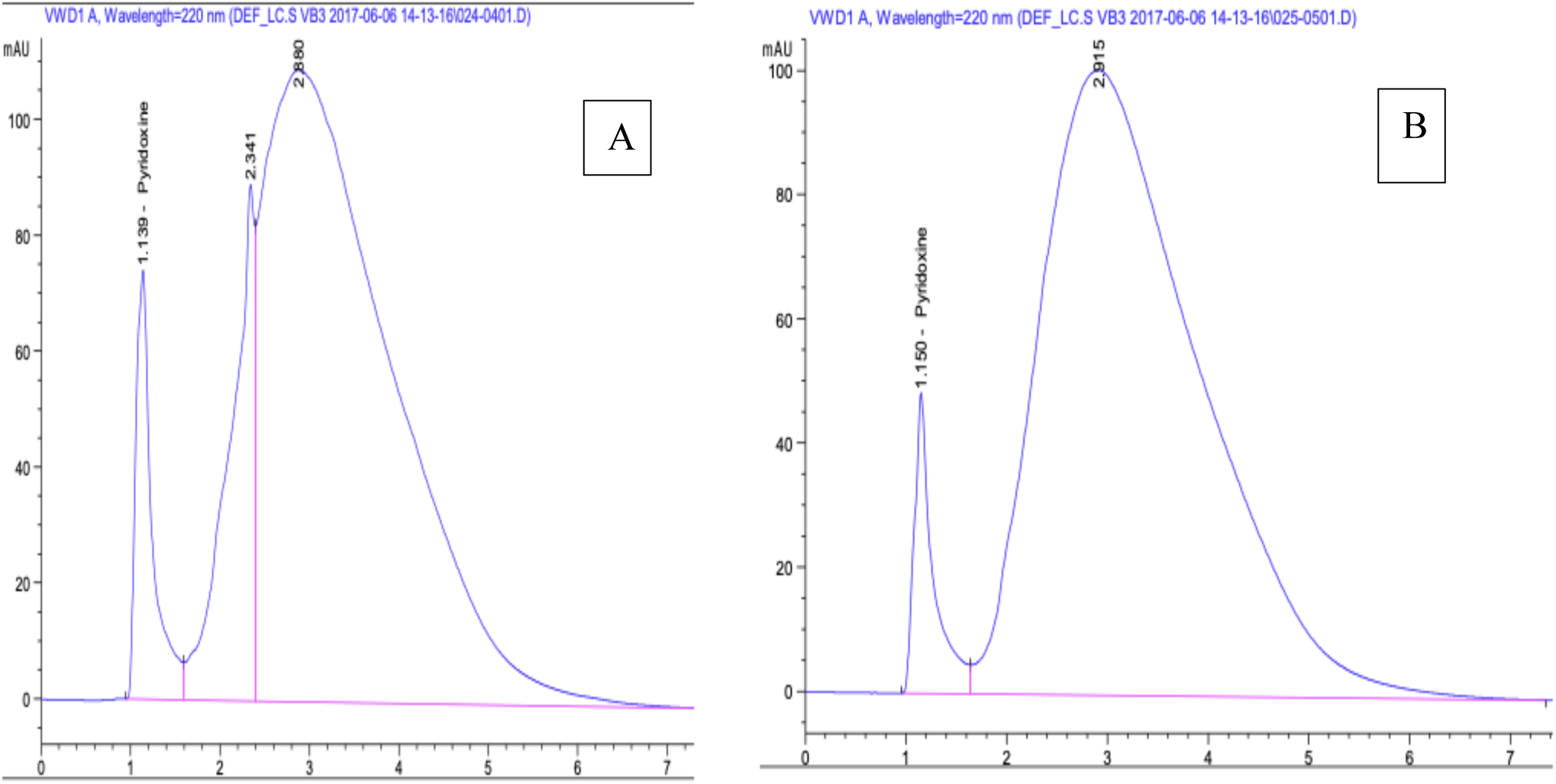
HPLC Spectra of Pyridoxine content of Site 1 (a) and 2 (b)

**Figure 3:**
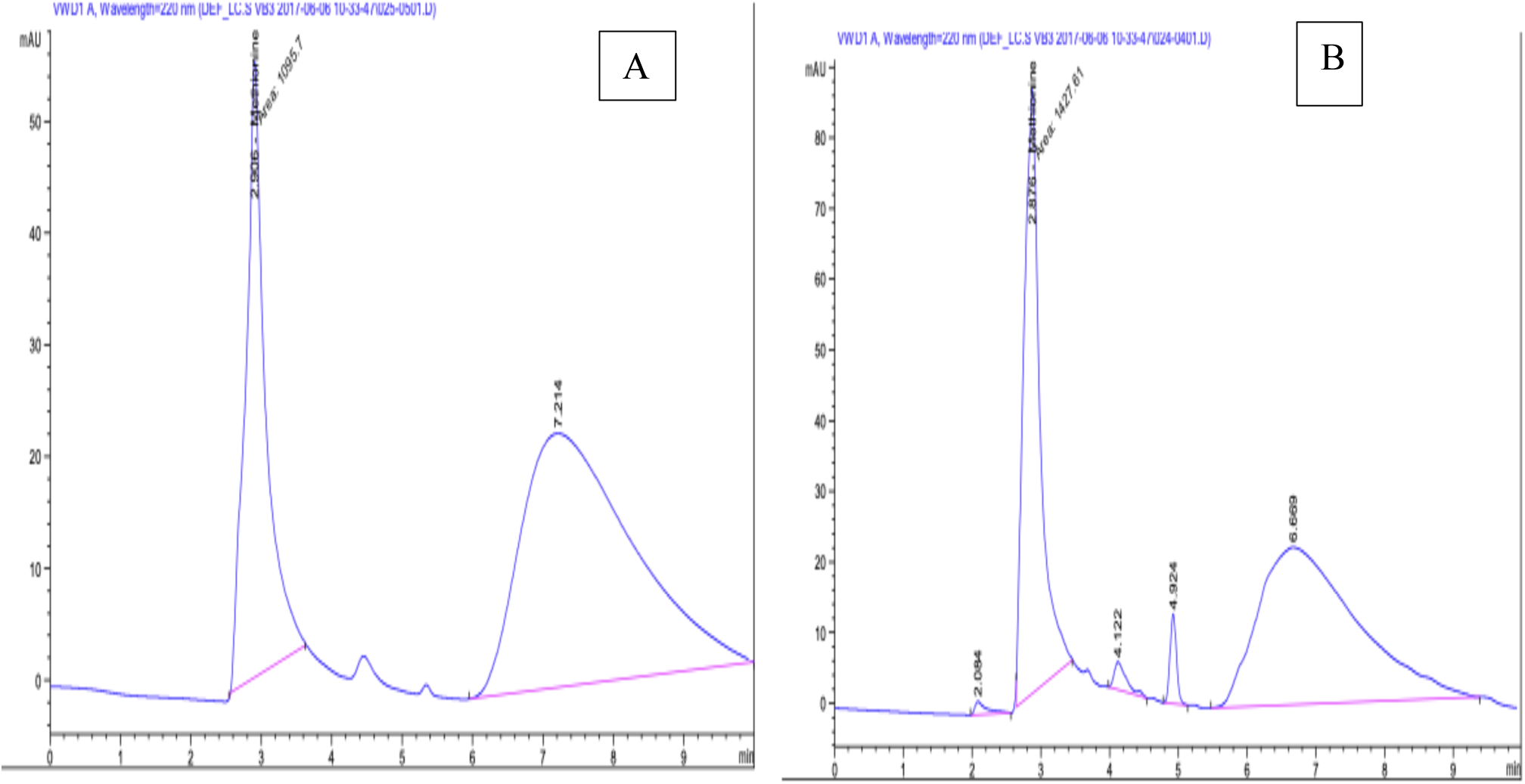
HPLC Spectra of Methionine Content of Site 1(a) and 2(b.

**Figure 4:**
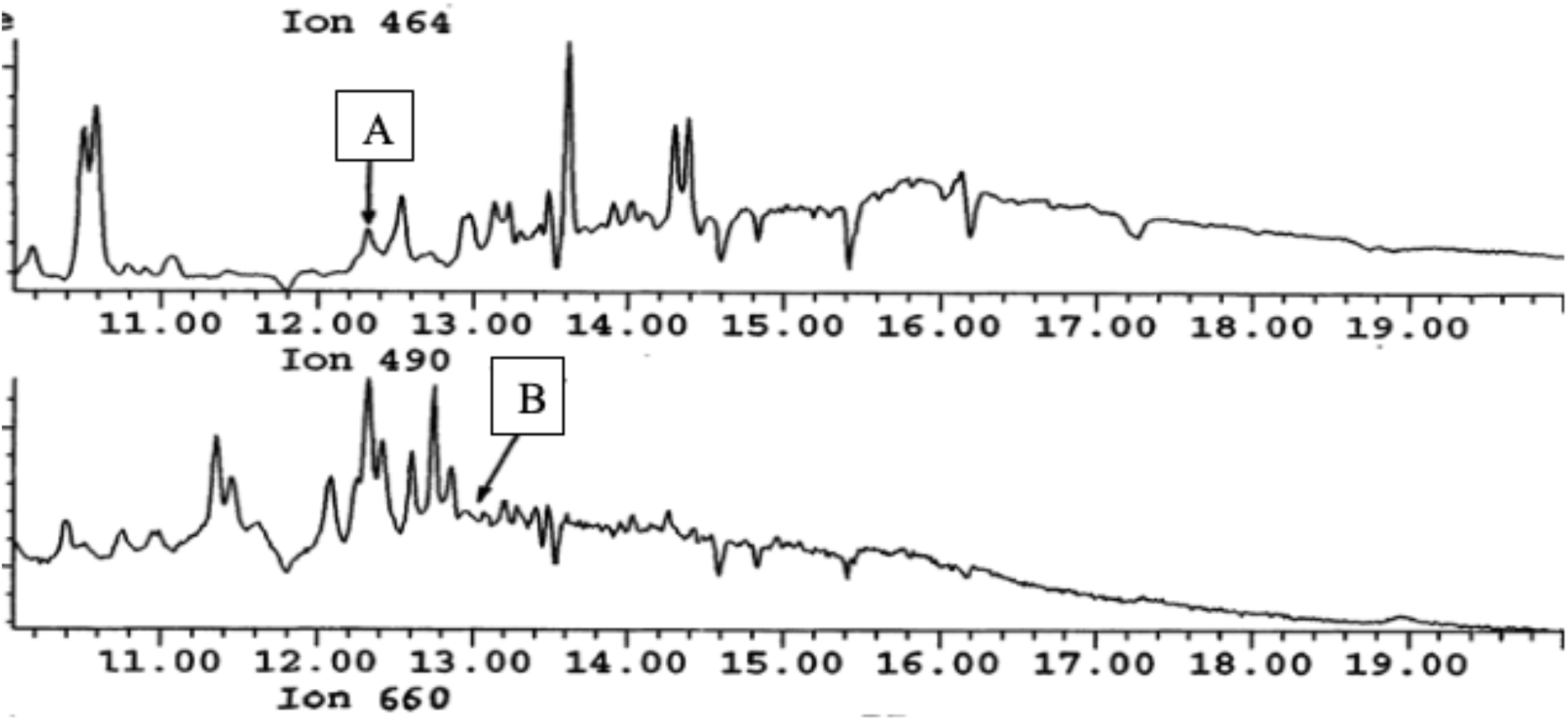
Chromatographic Spectra of estrone (a) and ethynylestradiol (b)

**Figure 5:**
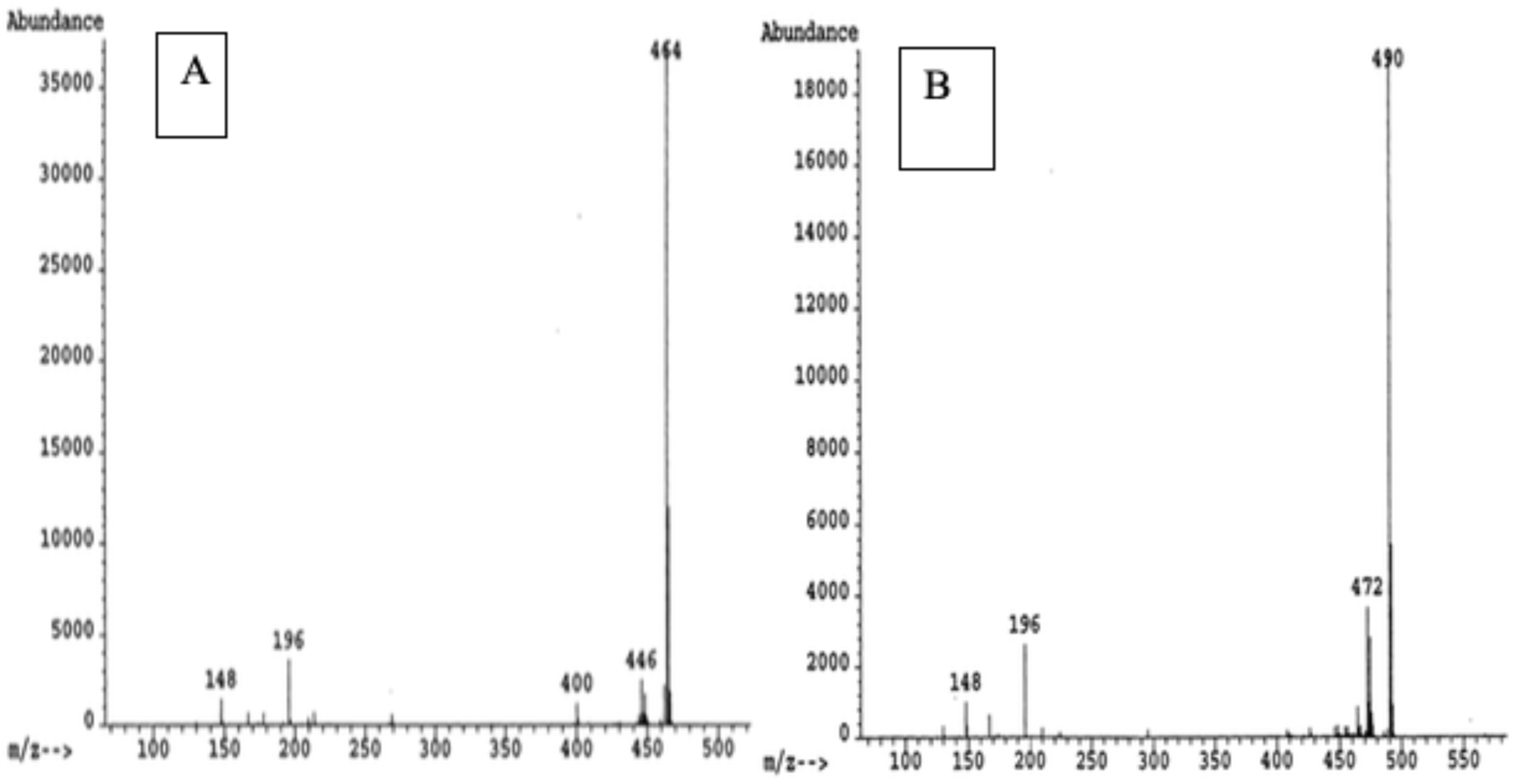
Mass Spectra derivative estrone (a) and ethynylestradiol (b)

### Collection of water samples

Water samples were collected from two locations, namely; in the conserved region of the Grove with limited human activity Site 1 (7°45’03.9”N and longitude of 4°33’03.9”E) and outside the Grove where there are unlimited activities, Site 2 (7°45’12.2”N 4°33’05.4”E) between April 2017 - September 2019 at 7am. Sample collection was subdivided to three, about 1000 mL each of water samples were collected in previously soaked with 10% HCL, washed with phosphate – free detergent, dried and pre – calibrated polythene screw capped plastic bottles. The remaining two portions were collected in clean high-density polyethylene (HDPE) dark bottles for vitamins analysis, amino acid assay as well as hormone content. All collected samples were immediately taken to the laboratory for analysis.

### Sample Analysis

#### Macroscopic Analysis of water samples

This was performed using the virtual and sensory evaluation method (Sharif *et al*., 2017). The colour, odour and the presence of foreign matters were observed.

#### pH determination

pH values of water samples were determined as described by (Raphael and Emmanuel, 2019). Prior to analysis, acidic and alkaline buffer solutions of pH 4 and 7 were used for calibration, readings were taken and water samples with pH values of less than 7 were considered acidic, pH = 7; neutral and greater than 7, alkaline.

#### Metal Analysis

Water samples was filtered through a 0.22 µm polypropylene Calyx capsule filter and collected in low-density polyethylene (LDPE) bottles. Sample was further acidified to pH < 2 with ultrapure grade hydrochloric acid (HCl) and stored cool for at least one month before extraction (Radulescu *et al*., (2014). Afterwards, samples were analysed using Atomic Absorption Spectrophotometer (AAS) as described by Smith, (1983).

#### Vitamins and Methionine Analysis

Vitamins and Methionine analysis were analysed using the liquid chromatographic method (Abano *et al*., (2014) and Cortés-Herrera *et al*., (2019). Water samples for vitamin and methionine analysis were filtered through 0.22µm polypropylene Calyx capsule filters and collected in high-density polyethylene (HDPE) dark bottles and stored frozen until analysis. Dissolved B-vitamins and methionine were extracted and pre-concentrated in solid-phase extraction onto a C_18_ resin before analysis.

#### Nutrient Analysis

Phosphate and nitrate analysis were performed according to the protocol described by Environmental Protection Agency, (2006).

##### Phosphate Analysis

Standard solutions were prepared by accurately measuring 10 mL of the stock solution into a 250 mL volumetric flask and made up to volume with distilled H_2_O. Varying volumes of the standard were then measured (5 mL, 10 mL, 15 mL, 20 mL and 25 mL) into separate labeled 100 mL volumetric flasks. The test water sample was diluted by a factor 10, before 25 mL of diluted sample was been transferred to a 100 mL volumetric flask, then made to mark using dilute distilled water. All solutions were kept for 30 minutes to allow color development before reading absorbance at 880 nm. Concentrations of the test samples were calculated from the standard curve.

##### Nitrate Analysis

Standard solutions were prepared by measuring 2 mL of the stock solution and made up to a 100 mL with distilled water. Varying volumes of the standard were measured into as separate beaker then interfering organic and metallic substances were removed by treating with 20 mL mercury (II) chloride solution. Two different volumes of each test sample were also subjected to similar treatment. All sample pH was adjusted to 11 with 50 % sodium hydroxide (NaOH), filtered to removed insoluble pellet. The initial flow through was discarded before allowing complete filtration. Then 2 mL of each filtrate was transferred into a beaker, and 1 mL of 1 % sodium salicylate solution was added, mixed well, and left to evaporate to dryness. It was later dried in the oven for 20 minutes at 105°C.

After removal from the oven, it was cooled, and then dissolved with 2mL concentrated tetraoxosulphate (VI) acid (H_2_SO_4_), 15 mL distilled water was added after the solution had cooled to room temperature followed by addition of 15mL of the sodium hydroxide – potassium sodium tartrate. The mixture was allowed to stand at room temperature for one hour and absorbance read at 420 nm.

##### Oestrogen Analysis

River water samples were prepared as described by Xiao *et al* 2001 using 8 ng/L estradiol II as internal control in each calibrated sample. The samples were then subjected to GCMS using the spitless technique, using 0.75 min period on an HP-5MS capillary column (15 m x 0.25 mm I.D., 0.25 mm film thickness) and 5% diphenyl – 95% dimethyl siloxane liquid phase. The oven temperature was maintained at 65°C for 1 min and then programmed to 220°C at 40°C per min, then to 255°C at 5°C per min and finally to 330°C at 20°C per min and maintained at 330°C. The injector and transfer lines were 330°C. Methane (99.99 %) was used as the reagent gas in the negative ion mode with source pressure of 160 Pa.

## 3. Result

### Sampling Area

Site one which is the major spiritual site of the grove with high natural conserved habitant, natural conservation, while site two is the area under the suspended bridge with minimal conservation as well as increased human and animal activities. Generally, the water body was free of macro – aquatic plantlike organisms and free flowing.

Table 1 shows the macroscopic (color, odour and foreign matter) characteristics and pH values for water samples collected from both sites. Both samples had similar characteristics with site 1 having a slightly high alkaline pH.

**Table 1:**
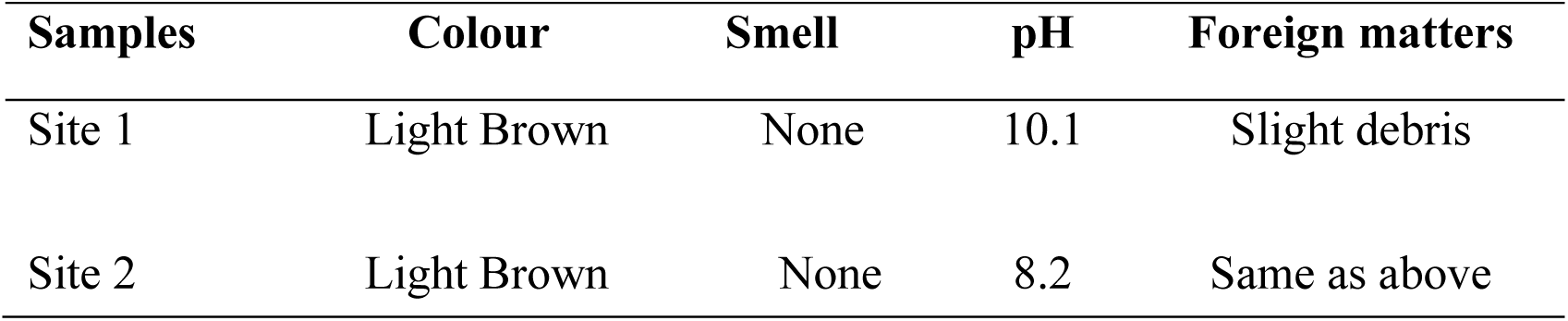
Macroscopic characteristics and pH of water samples from both sites.

| Samples | Colour | Smell | pH | Foreign matters |
| --- | --- | --- | --- | --- |
| Site 1 | Light Brown | None | 10.1 | Slight debris |
| Site 2 | Light Brown | None | 8.2 | Same as above |

### Sample Analysis

#### Metal Analysis

Plate I show a screenshot of the result of metal analysis, it showed that the average zinc content in site one was 0.079 mg/Kg, while that of site two was below detection. The manganese (Mn) content was practically the same for both sites, while the Nickel (Ni), Cobalt (Co) and Aluminum (Al) levels were almost the same throughout the study period for both sites.

#### Vitamins and Methionine Analysis

Vitamins and methionine content of the river water is represented (Table 1), the average Methionine content at site one was 74.41 ug/mL while site two was 57.11 ug/mL. The mean values of two water-soluble vitamins; Thiamine (vitamin B1) content of site one was 3.758 mg/Kg and 2.355 mg/Kg at site two and B6 (Pyridoxine) was 0.108 mg/Kg in site one and 0.072 mg/Kg at site two.

#### Nutrient Analysis

Phosphate and nitrate analysis done on the water sample yielded relatively lower concentrations in the water as shown in Plate 1. Average phosphate content of site 1 (s1) was observed to be 0.027mg/Kg while Nitrate content was 0.082mg/Kg.

#### Oestrogen Content

Over the stretch of the study period, the hormone values declined during the raining session by half from their maximum values for testosterone (4.8 ng/L), estrone (8.8 ng/L), ethinylestradiol (6.1 ng/L), and estrogen (4.9 ng/L) in site 1 estrogen (4.8 ng/L) and ethinylestradiol (2.4 ng/L) while estrogen was about ≥ 1.5 ng/L in site 2.

Over the same stretch, the hormone values declined by half from their maximum mean values for testosterone (3.3 ng/L), estriol (8.8 ng/L), ethinylestradiol (6.1 ng/L), and estrogen (4.9 ng/L). From 67 to 100 km mark, testosterone (4.8 ng/L) and estrogen (2.4 ng/L) were still elevated while ethinylestradiol and estriol were ≥ 1.5 ng/L.

## 4. Discussion

The potential contributions of drinking water to nutritional status also depend on water consumption, which is highly variable depending on both behavioral factors, believe, climatic as well as environmental conditions. Individuals with relative water consumption rate include infants, residents of hot climate, individual physiologically assumed to be susceptible to dehydration and individuals engaged in strenuous physical activity. The Osun Osogbo Heritage Grove is a major site of tourist attraction due to various beliefs that the water from the river possess the ability to heal and aid fertility. These beliefs led to thousands of devotees use the Osun river water for their daily needs such as drinking and bathing. Despite the growing concern on environmental issues and the increasing efforts on environmental monitoring and research, information on both river water composition and its quantity are fragmented and mostly gathered in unpublished reports. Thus, this research was carried out to give a background scientific knowledge on the constituents of the river, which are likely to aid understanding the role of some of these constituents in the acclaimed properties of the river water. Principal investigation of the micronutrient was based on their role in metabolism and in fertility (UNICEF/UNU/WHO 1999).

The results from this study revealed that the Osun river water is slightly brownish and highly alkaline pH. This slightly brownish might be attributed to the environmental pressure due to human activities from settlements along the river and rituals performed during festival that attract thousands of people (NCMM, 2005) and other anthropogenic factors which affect the properties of the water (Khatri and Tyagi 2015). While the alkaline pH = 10.1 of site 1 is higher when compared with Zamzam water with pH 8 (Shomar, 2012) and Mediterranean Sea water pH 8 (Flecha, 2015). Alkaline water are rich in minerals and attributed with health benefits such as ability to balance body pH, antioxidant, detoxification properties and generally optimised body immunity (Mousa, 2017). This could be attributed to the therapeutic value of the Osun River.

The presence of vitamins in drinking water has been of particular interest due to the role vitamins play in metabolism, especially the vitamin B complex family known to play significant role as cofactor in enzyme catalyzed reaction such as dehydrogenase complexes (Parra *et al*.,2018). Prominent among these vitamins are thiamine used in the synthesis of the cofactor Thiamine pyprophosphostate, pyridoxine and its role in the glycogen synthesis pathway as well as amino acid metabolism. In this study, the vitamins and methionine concentrations along the Osun river follow different trend, for instance, site one was observed to be richer in methionine (74.410 g/Kg), thiamine (3.75823 g/Kg) and pyridoxine (0.108020 g/Kg; 0.622776 g/Kg) when compared with site two where methionine (54.11 g/Kg), thiamine (2.35473 g/Kg), pyridoxine (0.0715691 g/Kg) values were detected respectively.

Conversely, an increase of vitamin B1 and B6 is observed in site one, when compared with site two, however, the values were lower than those reported for Moulouya river by Tovar-Sanchez *et al*. (2016), other vitamins such as B12 were not detected in the water samples. Opposite responses in the various B-vitamins is not rare since their availability in water is governed by the specificity of the predominant phytoplankton species for those vitamins (Sanudo-Wilhelmy *et al*. 2012). In this study, different values of vitamins (i.e., B1 and B6) were observed in the main worship area where the phytoplankton assemblages changed from dominance of diatoms to dinoflagellates mainly due to the fact that devotees tend to continuously drop sacrifice at this portion of the river. These might also give basis for the consistence slight brown coloration of the Osun water, going by the ability of dinoflagellates to generate “red tides”. In their report, Radi, *et al*. (2007) established the relationship between dinoflagellate cyst assemblages and hydrographic conditions, productivity and nutrient concentrations, they suggested that dinoflagellate cyst assemblages can be used to reconstruct primary productivity, temperature and salinity. Sa∼nudo-Wilhelmy *et al*. (2014) emphasized the regulatory role of Vitamins in metabolic activities of marine plankton. Because of their high bacterial activities, freshwater sources (such as rivers and groundwater) are considered important sources of vitamin B1 and B6 (Barada *et al*., 2013; Gobler *et al*., 2007; Okbamichael and Sa∼nudo-Wilhelmy, 2005).

The National *Agricultural Library* reported the role of trace metals such as: zinc, copper, manganese, etc. in the influence on reproduction and development. In a similar report by (Rasheed *et al*., 2003), NO_3_^-^ and PO_4_^3-^ were reported to play an important role in biochemical processes. Looking at the trace metal zinc, the value 0.079 g/Kg was obtained for site one, and - 0.015g/Kg for site two. Zinc, an essential metal which is needed for hormone regulation, immune builder and fertility in women was detected in the river sample at 7mg in each litre of water taken from the river, compared with standard FDA value of (3-5) mg/L. Aluminum content was observed to be 0.179g/Kg, site one and 0.192 g/Kg for site two, compared with standard FDA value of 0.05-0.2mg/L; this implies that for every litre of Osun water taken, 0.2 mg of aluminum is contained in it. The concentration of Cr in surface water represents the industrial activity (Shiller and Boyle, 1987). Surface water contains chromium in the range of 0.004 to 0.007 mg/L (Batayneh, 2012). The chromium, cadmium, copper and lead levels in the Osun River water were below detection indicating that the water from the River has zero concentration. Manganese (Mn) is the essential component of those biochemical reactions that affects bone, cartilage, brain and energy supply but toxic in higher concentration. Manganese in freshwater ranges from 1 to 200 μg/L. In this present study, the concentration of Mn determined was 0.313 g/Kg for both sites (Plate I) and do not exceed the FAO/WHO permissible limits for drinking water. Arsenic was 0.842 g/Kg for site one, and 0.569 g/Kg for site two, compared with 7.29 g/L reported by Fahad *et al*. (2016) for Zamzam. Although arsenic may cause low birth weight and spontaneous abortion, long-term chronic health effects, such as skin disease, skin cancer, it was and is still applied for pharmaceutical and medical purposes in curing asthma and hematological illnesses. In their report, Stein and Tallman (2012) described the use of Arsenic trioxide (ATO) as a new era in chemotherapeutic of acute promyelocytic leukemia (APL). A growing body of literature demonstrates the feasibility and efficacy of ATO, usually given with ATRA, in the treatment of patients with newly diagnosed APL. However, he mentioned reports of potential unintended toxicities, which included impaired fertility in both men and women. Walsh in his (2003) Second edition textbook of Biopharmaceutical Biochemistry and Biotechnology also describe biologic agent as any other trivalent organic arsenic compound applicable to the prevention, cure or treatment of disease or conditions of human beings.

Copper, cadmium, and lead had relatively no value (−0.006 g/Kg) when tested for in the Osun water; knowing that lead is harmful to the body, it was satisfactory to know the lead content of the Osun water was below detectable level throughout the period of study. Phosphate (PO_4_^3-^) value observed from the Osun water did not exceed the stipulated standard of 0.02 g/Kg, as the value obtained was 0.027 g/Kg. The NO_3_^-^ value obtained was 0.082 g/Kg. In summary, it was observed that higher nutrients levels were obtained from the first site, which is within the grove and the believed center of most of the spiritual activities of the devotees, and this is due to the natural conservation present over the river. Occurrence of metals such as Cu, Zn and Fe in water is also of importance considering the role of metals as cofactors of enzymatic activities and protein structure. In natural surface waters, the concentration of zinc is usually below 0.010 mg/L, while in groundwater 0.010–0.040 mg/L (Nriagu 1980; Elinder 1986). Essential amino acids such as methionine found in some water bodies has be attributed to environment or climatic conditions of the water. Micronutrients indirectly serve as the catalyst to release the energy from the macronutrients.

After obtaining the values 74.410 µg/mL for the first site, and 57.110 µg/mL for the second site, and knowing that methionine is an essential amino acid required for initiation of protein synthesis. It was satisfactory to know the methionine content is high when compared with standard FDA value 56.6 µg/mL. This might imply that an individual taking Osun water takes in over 55 µg of water dissolved methionine per every mL of the water. Vitamin B1 (Thiamine) content gotten in site one was 3.758µg/mL and site two was 2.355 µg/mL compared with standard of 1.5mg/l. Hence, it shows that if one takes a mL of Osun water, the thiamine content obtained from it is over 3µg compared with the RDA value of 1.1mg. Vitamin B6 (Pyridoxine) value obtained was 0.108 µg/mL for site one, while 0.072 µg/mL was observed for site two, and this shows that for every mL of the Osun water taken in, 0.1µg of pyridoxine is contained in it. Due to the high bacterial activities, freshwater sources (such as rivers and groundwater) are considered important sources of vitamin B_1_ and B_6_ and Baren-cohen *et al*, (2006) reported that hormones in readily measured quantities can be transported along a considerable distance from the source of pollution.

Several literatures have shown that steroid hormones produced by humans and animals constantly excreted into the environment found their ways into underground water and rivers (Lintelmann *et al*., 2003; Drewes and Shore, 2001; Shore and Shemesh, 2003). This work concentrated on naturally occurring hormones such as estrone (E_1_) and estradiol-17b (E_2_) which were reported to exert physiological effect at concentrations above LOEL (Lowest observable effect level). E_2_ is abiotically converted to E_1_ thus, they are generally considered as oestrogen. The LOEL for E_2_ and E_1_ were report as 14 and 3.3 ng/L, respectively (Olsen *et al*., 2007) while ethinylestradiol is 1 ng/L (Baren-cohen *et al*, 2006). The mean values of steroid detected in the Osun River water over the study period shows the hormone content were lower doing pre-raining season, but the content was both above the LOEL. Ethinyl estradiol binds to the estrogen receptor complex and enters the nucleus, activating DNA transcription of genes involved in estrogenic cellular responses. This agent also inhibits 5-alpha reductase in epididymal tissue, which lowers testosterone levels and may delay progression of prostatic cancer. In addition to its antineoplastic effects, ethinyl estradiol protects against osteoporosis. In animal models, short-term therapy with this agent has been shown to provide long-term protection against breast cancer, mimicking the antitumor effects of pregnancy

In conclusion, this study established micronutrient, trace metals, water soluble vitamin, hormone content as well as other dissolved trace metals, Pb, Cd, As, Zn, Cu, Ni, Co, Fe, Mn, Cr and PO^4^ which were within the acceptable limits. Hence, maybe associated with metabolic and physiological processes leading to the therapeutic claims of the ancient River.

## Supporting information

cover letter

