## Supplementary material for "Nutrients: trace metals, micronutrients, oestrogen and B-vitamin content of Osun River: A River that runs southwestern Nigeria into the Atlantic Gulf of Guinea": cover letter

**FOUNTAIN UNIVERSITY, OSOGBO**
(UNIVERSITY OF THE NASRUL-LAHI-IL-FATIH SOCIETY OF NIGERIA)
COLLEGE OF NATURAL AND APPLIED SCIENCES
DEPARTMENT OF CHEMICAL SCIENCES (BIOCHEMISTRY UNIT)
**Molecular Biology & Genetic Diversity Research Laboratory**

20th August 2020.

The Editor-in-Chief
PLOS Biology

**SUBMISSION OF MANUSCRIPT.**

On behalf of my co-author, I write to submit a manuscript tiled "**Nutrients: trace metals, micronutrients, oestrogen and B-vitamin content of Osun River: A River that runs southwestern Nigeria into the Atlantic Gulf of Guinea**" being finding from our research on the Osun River water constituents in a bid to provide a scientific insight for the constituent of the widely used river water.

I look forward to receiving any comments that you may have in relation to the above. You can contact me via

Thanking you in anticipation of a favourable consideration.

Yours sincerely

Afolabi-Balogun Nusrah Bolatito
(Corresponding Investigator)

FUO_MBGE_RG
